# *dach-1*, a cytochrome P450 gene, regulates synaptic and developmental plasticity during the diapause of *Caenorhabditis elegans*

**DOI:** 10.1101/2022.07.05.498867

**Authors:** Sangwon Son, Myung-Kyu Choi, Daisy S. Lim, Jaegal Shim, Junho Lee

**Affiliations:** Department of Biological Sciences, Institute of Molecular Biology and Genetics, Seoul National University, Seoul, 08826, Korea; Research Institute, National Cancer Center, Goyang, 10408, Gyeonggi, Korea

**Keywords:** phenotypic plasticity, dauer, aldicarb, synaptic transmission, dauer-specific acetylcholine defect -1 (*dach-1)*, cytochrome P450, *cyp-34A1*

## Abstract

Animals exhibit phenotypic plasticity through the interaction of genes with the environment, and little is known about the genetic factors that change synaptic function at different developmental stages. Here, we investigated the genetic determinants of how developmental stages alter synaptic transmission using the free-living nematode *Caenorhabditis elegans. C. elegans* enters the stress-resistant dauer larval stage under harsh conditions. Although dauer is known to have reduced permeability and increased resistance to most known exogenous chemicals, we discovered that dauer is hypersensitive to a cholinesterase inhibitor, aldicarb. To investigate genes regulating dauer-specific acetylcholine transduction, we first screened for aldicarb-resistant mutations in dauer and then performed a secondary screen to rule out aldicarb-resistant mutations that also affect adults. We isolated two different mutations of a single gene called *cyp-34A4* or *dach-1* encoding a cytochrome P450. In the non-dauer stages, *dach-1* is mainly expressed in the intestine, but its expression is robustly increased in the epidermis of dauers. By tissue-specific rescue experiments, we found that *dach-1* modulates aldicarb sensitivity in a cell non-autonomous manner. In addition, *dach-1* plays pleiotropic functions in dauers by regulating quiescence and surviving heat shock and hyperosmolar stress. Our study reveals novel functions of the cytochrome P450 in synaptic and physiological changes during developmental plasticity.

## Introduction

Various molecular mechanisms of synaptic transmission play their roles at specific times and specific anatomical locations. There are spatially distinct mechanisms such as neurotransmitter release in the presynapse region and receptor levels in the postsynaptic region. Released neurotransmitters are regulated through degradation or reuptake. Furthermore, neuronal activity is regulated by non-neuronal tissues such as glia. For example, a prostaglandin released from astrocytes regulates the excitability of GnRH neurons through paracrine signaling (Clasadonte *et al*. 2011) and thus is called a gliotransmitter important for glia-to-neuron communication.

To understand hidden mechanisms of synaptic dysfunction and find novel therapy targets, non-biased screening is one of the most efficient strategies. Pharmaceutical treatment has been widely used to investigate synaptic transmission combined with non-biased screening (Jones *et al*. 2005). One of the most commonly used chemicals in the model animal *C. elegans* is an acetylcholinesterase inhibitor called aldicarb (Blazie and Jin 2018). A working concentration of aldicarb increases the amount of acetylcholine at the neuromuscular junction (NMJ), causing paralysis of the animals within a few hours (Mahoney *et al*. 2006; Oh and Kim 2017). In this condition, mutations affecting synaptic transmission change the time course of aldicarb-induced paralysis. Such Ric (Resistance to an inhibitor of cholinesterase) phenotype causing slow aldicarb-induced paralysis is associated with reduced acetylcholine synthesis, vesicle loading, vesicle release, or reduced acetylcholine receptor (Nguyen *et al*. 1995; Rand 2007). Although the Ric phenotype was due to the over-accumulation of acetylcholine, the revealed functions of most Ric genes are not limited to acetylcholine transmission, but rather are involved in all neurotransmitters. This is because vesicle loading or release mechanisms are usually not specific to a particular neurotransmitter. For this reason, the Ric phenotype is an efficient method to find novel genes important for the overall function of synaptic transmission. Although Ric genes were found by many studies, all studies were focused on the phenotype in adults, and no studies were conducted in the different developmental stages.

The onset of neuronal dysfunction often depends on certain conditions such as the developmental stage (Zhang *et al*. 2017). It is because mechanisms regulating synaptic activity before adulthood are different from those after reaching the adulthood. For example, in the human adolescent brain, the activity of the brain region related to reward is relatively elevated, whereas the region related to the aversive stimulus is relatively weakened (Spear 2013). In the case of *C. elegans*, new postembryonic motor neurons are differentiated during the transition from L1, the first larva period, to the next developmental stage, and the position of the synapse of the pre-existing motor neurons is remodeled (Rapti 2020). Structural changes of neurons according to developmental stages suggest that synaptic transmission may also vary in specific developmental stages. If the synaptic transmission is increased in a certain stage, aldicarb-induced paralysis might be observed even at the sub-threshold level of the adult stage. In this study, we utilized the sub-threshold level of aldicarb to test whether other developmental stages elicit increased synaptic transmission. Then, we focused on the aldicarb hypersensitivity in the developmental stage called dauer. Based on the dauer-specific aldicarb hypersensitivity, we conducted a forward genetic screening to discover the mutations that induce dauer-specific aldicarb resistance. From the screening, we found that two alleles of *dach-1* or *cyp-34A4*, caused a dauer-specific Ric phenotype. *dach-1* has not been isolated from conventional Ric screening using adults and appears to be a mechanism to regulate synaptic transmission through paracrine factors from non-neuronal tissues. Furthermore, we discovered that *dach-1* is important for other phenotypic plasticity in dauer.

## Materials and methods

### Maintenance and strains

Most worms were maintained at 20°C as presviously described (Brenner 1974), and *daf-2(e1370)* and other *daf-2(e1370)* background mutants were maintained at 15°C. The following strains were used: N2, *daf-2(e1370), daf-2(e1370);dach-1(ys51), daf-2(e1370);dach-1(ys52), dach-1(ys51), dach-1(ys52), rrf-3(pk1426);daf-2(e1370), rrf-3(pk1426);daf-2(e1370);dach-1(ys51)*, N2;*Ex[Pdach-1::GFP, rol-6(su1006)], dach-1(ys51);Ex[Pdach-1::dach-1::SL2::GFP, Pmyo-2::mCherry], dach-1(ys51);Ex[Pdpy-7::dach-1::SL2::GFP, Pmyo-2::mCherry], dach-1(ys51);Ex[Pmyo-3::dach-1::SL2::GFP, Pmyo-2::mCherry], dach-1(ys51);Ex[Pegl-3::dach-1::SL2::GFP, Pmyo-2::mCherry], ace-3(dc2), ace-3(dc2);dach-1(ys51)*.

### Molecular biology

Each promoter was sub-cloned by restriction enzymes respectively. Unspliced genomic sequence of *dach-1* containing 3’ UTR was inserted into pPD114.108(SL2::GFP) vector using BSSHII and NotI as the restriction sites. *Pdach-1, Pdpy-7, Pmyo-3*, and *Pegl-3* were inserted into the *dach-1::SL2::GFP* vector using SalI and BSSHII as the restriction site. All the promoter information was gained from the Promoterome database (Dupuy *et al*. 2004).

### Generation of transgenic lines

Introducing DNA into the gonads of young adult hermaphrodites was carried out as previously described (Mello *et al*. 1991). After microinjection, worms were quickly recovered with M9 buffer. For the reporter transgenic lines, *rol-6(su1006)* was used as the injection marker with 50 ng/μl of concentration. For the rescue experiments, *Pmyo-2::mCherry* was used as the marker with 3 ng/μl concentration, along with 50 ng/μl of rescue construct and 47ng/μl of empty vector (pPD95.77).

### Ethanol and stress sensitivity assay

7% ethanol sensitivity assay was conducted as previously described (Choi *et al*. 2016). First, 50 worms were moved to an empty, unseeded NGM plate. Dauer larvae were transferred via mouth pipette with M9 buffer, and L3 or non-dauer larvae were done via Platinum pick. After 10 minutes, they were harvested with 1 ml solution containing 7% ethanol and immersed as a droplet in an empty 55 mm petri dish. The lid was closed and swimming worms were counted. For heat stress, non-seeded NGM plates containing about 20 dauers ware incubated at 37°C for 4 hours, then transferred to 20°C, and the number of live and dead insects was counted the next day. At a heat stress of 30°C, most of the dauers did not die, so 37°C was used as the heat stress condition (data not shown). For high osmotic stress, about 20 worms were transferred to a solution containing 1500 mM NaCl with a mouse pipette and incubated at 25°C for 20 hours. After washing and harvesting the worms, they were transferred to an NGM plate and the number of moving and dead worms was counted the next day.

### Dauer formation

Dauer formation was induced with the cocktail of ascr #1, #2, and #3 in NGM plate. For 250 ml of pheromone plates, 0.5 g NaCl, 0.75 g KH2PO4, 0.125 g K2HPO4, 5 g agar were dissolved in 248 ml DW and autoclaved. Before pouring 0.5 ml Cholesterol, 0.5 ml Daumone 1 (ascr #1 “C7”), 0.5 ml Daumone 2 (ascr #2 “C6”), and Daumone 3 (ascr #3 “C9”) were added (Ludewig *et al*. 2013). Bactotryptone was not included in the pheromone plates to limit the growth of bacteria. After one day, 100 μl of saturated OP50 culture was seeded and incubated in room temperature for another day, and stored at 4°C for up to 1 month. For inducing dauer formation, 7∼10 young adults were placed on a pheromone plate and incubated at 25°C for 4 days. For inducing dauer formation in the *daf-2(e1370)* background, worms were not placed in pheromone plates but NGM plates instead. Several L4 worms were moved to a new NGM plate and incubated at 15°C for 2 days. After L4 worms were grown into adults and laid enough numbers of eggs, the NGM plate was incubated at 25°C for 4 days.

### Quantification of nictation behavior

For quantifying nictation behavior, the protocol from (Lee *et al*. 2012) was followed. 50 dauer larvae were harvested with M9 buffer and transferred to a microdirt chip. After 30 minutes, the nictation ratio of each worm was tested for 1 minute, and the tested worm was eliminated from the microdirt chip. For each set of experiments, 15 worms were tested and this procedure was repeated 2 times to 3 times.

### Quantification of dauer quiescence

On the dauer-induced phermone plate, the total number of dauers and the number of moving dauers were counted. After three measurements on one plate, the average value was recorded as a representative value. Independent experiments were repeated three times.

### Feeding RNA interference method

Clones of C. elegans RNA interference library were from Ahringer or Vidal Library. All the RNAi clones were carried by HT115 bacterial cell line. Each RNAi cell was streaked and cultured on LB containing Ampicillin, and transcriptional activation was induced by 1 mM IPTG. *rrf-3(pk1426);daf-2(e1370)* double mutant was used to conduct RNAi screening in dauer stage. Worms were placed on RNAi plates at L4 stage and incubated at 15°C. After one generation, L4 stage worms were transferred to new RNAi plates. After 2 days, young adults with eggs were moved to 25°C, and dauer formation was induced. After 4 days dauer larvae outside the E. coli lawn were harvested and placed on aldicarb plates for the assay.

### Aldicarb assay

100 mM aldicarb stock was dissolved in 70% ethanol. For 250 ml of aldicarb plates, 0.5 g NaCl, 0.75 g KH2PO4, 0.125 g K2HPO4, and 5 g agar were dissolved in 250 ml DW and autoclaved. Before pouring, add 250 μl of 100 mM aldicarb stock (for a final concentration of 0.1 mM aldicarb). One day after pouring, 0.1 mM aldicarb plates were stored at 4°C for up to 1 month. For comparison of aldicarb sensitivity of dauer larvae, 10 worms were harvested with M9 buffer and placed on the 0.1 mM aldicarb plate. After 70 or 80 minutes, moving worms on the plate were counted. Every experiment was repeated 3 times.

### EMS mutagenesis

To isolate aldicarb-resistant dauer mutants, random mutagenesis was conducted using EMS. *daf-2(e1370)* mutant was used as wild type. Young adult worms of *daf-2(e1370)* were treated with a final concentration of 50 mM EMS for 4 hours and washed with M9 buffer. After EMS treatment, samples were treated with an inactivation solution containing sodium thiosulfate (806 mM) and NaOH (100 mM). Mutagenized P0 worms were placed on half-seeded NGM 100 mm plated. The total number of F1 progeny was about 37,800. When F1 worms lay eggs, they were moved to 25°C. After 4 days, dauer-induced F2 offspring worms were harvested with M9 buffer and placed on 0.1 mM aldicarb-containing 100 mm plates. After 80 minutes, each moving worm was selected and placed on NGM plates respectively and was incubated at 15°C for dauer recovery.

### Whole-genome sequencing

To identify the causal genes of *daf-2(e1370);ys51* and *daf-2(e1370);ys52*, whole-genome sequencing was conducted. *daf-2(e1370);ys51* and *daf-2(e1370);ys52* were outcrossed with *daf-2(e1370)* two times independently. F2 homozygote mutants were screened through the aldicarb-resistant phenotype as described above. At the second outcross, more than 10 independent homozygote mutants were isolated for each allele. Genomic DNA was extracted for each mutant, and the same amounts were mixed. Finally sequenced samples were *daf-2(e1370);ys51* and *daf-2(e1370);ys52*. The Illumina sequencing condition was HiSeq4000/ 100PE / 5Gb. For the process of raw data, MAQGene software and Wormbase WS195 reference genome were utilized (Bigelow *et al*. 2009). The final “grouped” information was exported to an excel file, and all the mutation information was grouped into chromosomes. For each allele, all the mutations shared with any other alleles were excluded, and allele-specific mutations were sorted. For each allele-specific mutation, the percent homozygosity was obtained by diving the number of variants read into the number of wildtype reads. Mutations with percent homozygosity over 95 were sorted. Then C→T or G→A substitutions which are usually induced by EMS mutagenesis, and missense or premature stop classes were finally sorted.

### Microscopy

Fluorescence images were taken through a confocal microscope (ZEISS LSM700, Carl Zeiss, Inc). Worms were harvested with 2.5 mM Levamisole (dissolved in M9 buffer) and placed on an agarose pad.

### Data Availability Statement

Strains and plasmids are available upon request. The authors affirm that all data necessary for confirming the conclusions of the article are present within the article, figures, and tables.

## Results

### Dauers are more sensitive to cholinesterase inhibitors, but not to an acetylcholine agonist

In adult worms, paralysis has been observed in 0.5 to 1.5 mM concentration of aldicarb (Mahoney *et al*. 2006; Oh and Kim 2017). However, 0.1 mM aldicarb induces uncoordinated movement but does not induce paralysis. We tested whether a sub-threshold, 0.1 mM aldicarb induced paralysis in other developmental stages and discovered that 0.1 mM aldicarb-induced paralysis was induced only in dauer, a hibernation stage after harsh conditions (Figure 1A and B). The dauer-specific aldicarb hypersensitivity was a surprising result because dauer is known to be impervious to many external chemicals. Indeed, we confirmed that dauer had more resistance than L3 to 7% ethanol, which induces paralysis by changing neuronal activity (Choi *et al*. 2016) (Figure 1C). We compared the responses of dauer larvae to another kind of cholinesterase inhibitor, trichlorfon. It is an organophosphate cholinesterase inhibitor with a different chemical structure from that of aldicarb (Nguyen *et al*. 1995). We confirmed that dauer larvae were sensitive to trichlorfon as well (Figure 1D), again showing increased cholinergic transmission in dauer.

**Figure 1.**
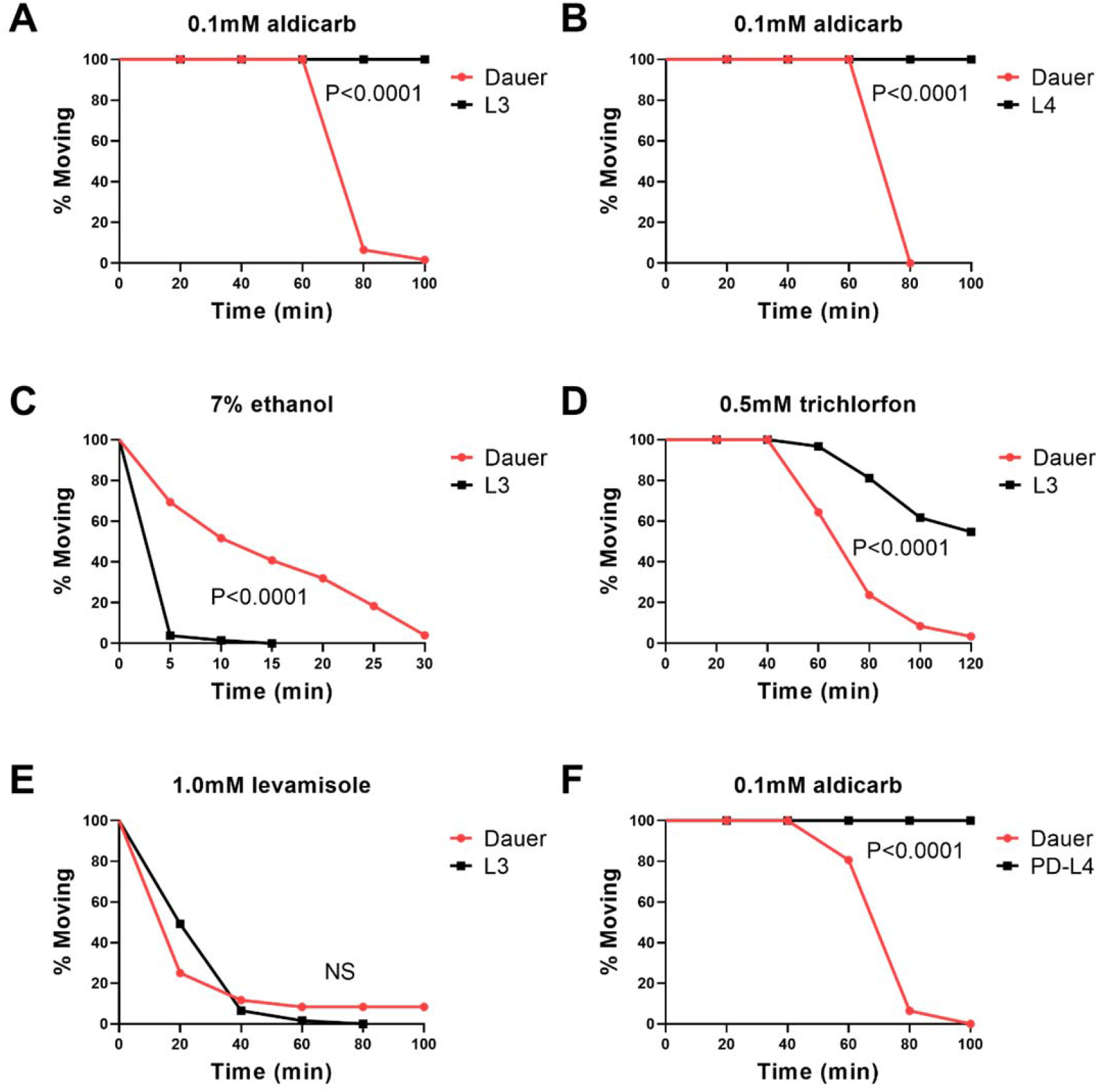
Altered acetylcholine transmission in dauer stage of *C. elegans*. (A, B) Dauer was sensitive to aldicarb compared to L3 stage (A) and L4 stage (B) larvae. (Band trichlorfon, cholinesterase inhibitors, compared to L3 stage larvae. (C) Dauer was resistant to 7% ethanol compared to L3. (D) Dauer was sensitive to trichlorfon compared to L3. (E) Dauer was not sensitive to levamisole compared to L3. (F) Postdauer-L4 was not sensitive compared to dauer. Statistical analysis is perfomed by Log-rank (Mantel-Cox) test test.

Paralysis due to excess accumulation of acetylcholine occurs primarily through levamisole-sensitive nicotinic Ach receptors expressed in muscle (Rand 2007; Treinin and Jin 2021). In contrast to dauer-specific aldicarb hypersensitivity, dauer did not show significant hypersensitivity upon levamisole compared to L3, and some dauers were still moving even when L3 showed complete paralysis (Figure 1E). It suggests that dauer-specific aldicarb hypersensitivity is not due to postsynaptic mechanisms. Next, we investigated whether changes in synaptic transmission in dauer were maintained during subsequent developmental stages. When a dauer encounters a favorable condition, it develops into an adult after passing through postdauer-L4 (PD-L4). Unlike dauer, PD-L4 had aldicarb resistance similar to that of normally developed L4 (Figure 1F). It demonstrates that the increased synaptic transmission in dauer is not imprinted from the experience of early development, but is rapidly changed by adapting to a new environment.

### *dach-1*, a cytochrome P450 gene, regulates neurotransmission in the dauer stage

To elucidate the mechanism of developmental stage-specific alteration of acetylcholine transmission, we performed a forward genetic screening for dauer-specific aldicarb-resistant mutants (Figure 2A). To induce dauer formation on large scale without dauer-inducing pheromone, a dauer-constitutive mutant, *daf-2(e1370)* was used for mutagenesis (Fielenbach and Antebi 2008). *daf-2* is the insulin/IGF receptor homolog and the mutation induces dauer at the restrictive temperature (Kenyon *et al*. 1993; Kimura *et al*. 1997). Similar to N2 dauer, *daf-2(e1370)* dauer was sensitive to aldicarb compared to L3 (Figure S1A). We conducted EMS mutagenesis following the standard protocol (Brenner 1974). The number of F1 progeny from 420 P0 worms was 37,800 when laying enough F2 eggs at permissive temperature. Then we induced dauer by transferring the F2 eggs at the restrictive temperature, then screened aldicarb-resistant F2 dauer (Figure 2A). After discarding the aldicarb resistant mutants that show phenotype both in dauer and adult, two independent mutants that showed phenotype only in dauer were isolated – *ys51* and *ys52* (Figure 2B and Figure S1B). Each mutant was outcrossed with *daf-2(e1370)*, and whole-genome sequencing was conducted to identify the causal gene for each mutant. Genomic DNA from pooled recombinants was sequenced, and homozygous mutations were sorted and analyzed (Doitsidou *et al*. 2010) (Figure S2). As the result, we have discovered that *ys51* and *ys52* shared distinctive premature stop mutations at the different locations of T09H2.1, a cytochrome P450 gene *cyp-34A4* (Figure 2C). *ys51* and *ys52* failed to complement each other, confirming that they are two different alleles of the T09H2.1 mutant (Figure S3). Also, we found that knockdown of T09H2.1 with feeding RNAi showed the aldicarb resistance in the *rrf-3(pk1426);daf-2(e1370)* background (Figure 2D). Taking these results together, we concluded that *cyp-34A4* regulates dauer-specific alteration of acetylcholine transmission, and named the gene *dach-1* (<u>d</u>auer-specific <u>Ach</u> transmission defect-1).

**Figure 2.**
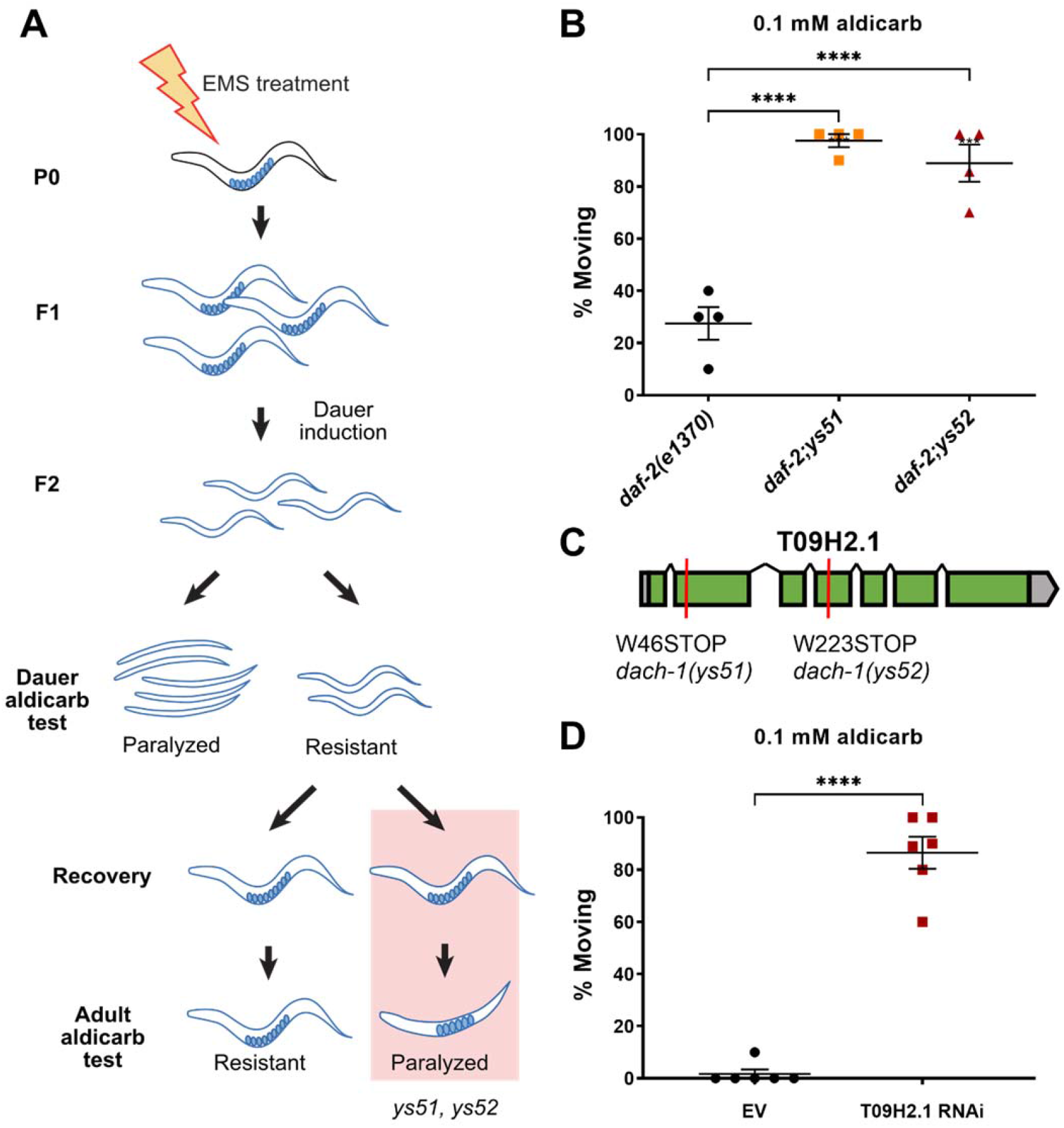
*dach-1 is* the cytochrome P450 regulating acetylcholine transmission in the dauer stage. (A) Experimental scheme of forward genetics for dauer-specific aldicarb-resistance. (B) *ys51, ys52* mutants showed aldicarb-resistant phenotype in the dauer stage. **** P<0.0001 (One-way ANOVA, Dunnett’s post-test). (C) *ys51* and *ys52* forms distinctive premature stop codon in T09H2.1 (D) Feeding RNAi of T09H2.1 in a RNAi-sensitive sensitive *rrf-3;daf-2* phenocopied aldicarb-resistant phenotype in the dauer stage. **** P<0.0001 (Unpaired t-test). For (B) and (D), aldicarb sensitivity was measured as the ratio of paralysis at 80 min.

### *dach-1* is expressed in epidermis from dauer and can exhibit cell non-autonomous effects

To elucidate the role of *dach-1*, we analyzed its spatiotemporal expression. The transcriptional reporter of the 1.2 kb sized *dach-1* promoter with GFP is only expressed in the intestine from the non-dauer, which is consistent with the previously reported expression pattern of the 2.0 kb sized *dach-1 promoter* (Joshi *et al*. 2021) (Figure 3A). During dauer, the transcription of *dach-1* showed a robust increase in non-intestinal tissues (Figure 3B). We confirmed that the epidermis is the tissue in which *dach-1* expression is specifically upregulated in dauers by colocalization with *Pdpy-7*, an epidermis-specific promoter (Figure 3C). The aldicarb-resistance phenotype of the *dach-1* mutant was rescued when the wild-type cDNA of *dach-1* was expressed in the mutant using the same promoter used in transcriptional reporter (Figure 3D). Consistently, we confirmed that rescue was successfully achieved when *dach-1* cDNA was expressed using an epidermis-specific promoter (*Pdpy-7*). Furthermore, rescue experiments were also performed by ectopic expression of *dach-1* cDNA in body wall muscle (*Pmyo-3*) or the nervous system (*Pegl-3*). Although the *dach-1* transcriptional reporter was not expressed in neurons or muscles, the ectopic expression of *dach-1* cDNA in muscles or neurons sufficiently rescued *dach-1* mutant. Next, we tested whether the role of *dach-1* in aldicarb treatment is truly mediated by the level of endogenous acetylcholine which is released onto NMJ. Released acetylcholine is normally degraded by cholinesterase. If cholinesterase is mutated, the level of extrasynaptic acetylcholine is increased. We confirmed that *ace-3*, one of cholinesterase mutants suppresses the aldicarb resistant phenotype of *dach-1* (Rand 2007; Treinin and Jin 2021) (Figure 3C). Overall, the results indicate that although *dach-1* is not expressed in neurons or muscles, its cell non-autonomous effect can regulate synaptic transmission in NMJ. *dach-1* encodes a cytochrome P450 enzyme, therefore, its enzyme products are expected to play a role in the paracrine effect regulating the synaptic transmission of NMJ from the epidermis. The rescue effect of *dach-1* by ectopic expresion in the neuron and muscle suggests that the substrate of *dach-1* may exist in other tissues as well as in the epidermis. Another *C. elegans* cytochrome P450 gene, *daf-9*, synthesizes a steroid hormone called dafachronic acid from cholesterol, which inhibits dauer entry (Gerisch *et al*. 2001). Accordingly, when cholesterol depletion occurs, dauer entry increases because DAF-9 cannot synthesize dafachronic acid. We investigated whether *dach-1*, similar to *daf-9*, is involved in the synthesis of cholseterol-derived steroid hormone and modulates aldicarb sensitivity. We found that the aldicarb sensitivity of dauer did not change under the condition of cholesterol depletion (Figure S4). Thus, *dach-1* appears to regulate neurotransmission through mechanisms other than steroid hormone metabolism.

**Figure 3.**
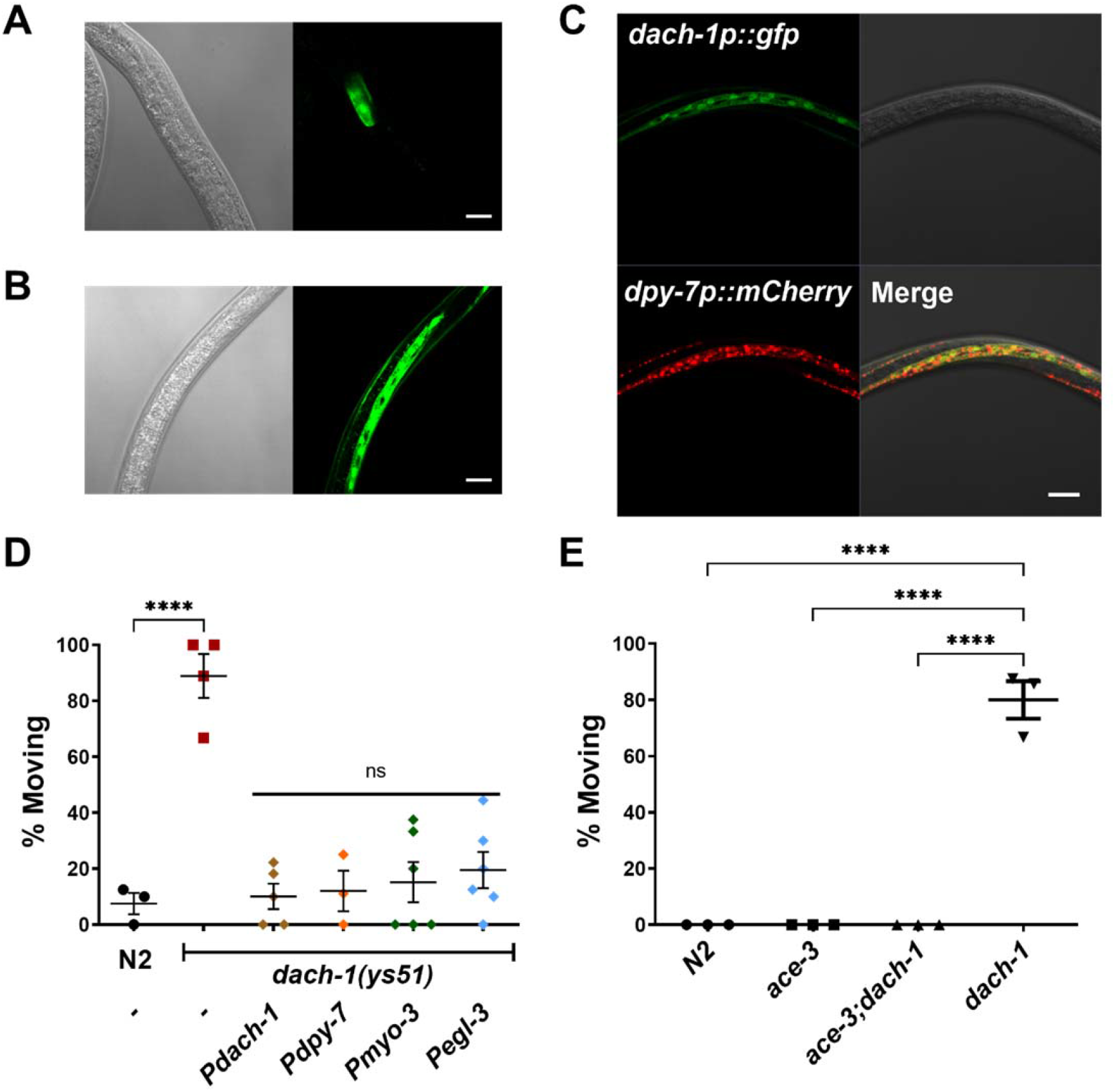
*dach-1* is increased in epidermis in dauer stage and acts in cell non-autonomous manner. (A) 864 bp upstream from start codon of *dach-1* was labeled with GFP. Expression of *dach-1* was shown in intestine at L4 stage. Bar, 20 μm. (B) *dach-1* promoter showed increased expression at dauer stage. Bar, 20 μm. (C) *dach-1* promoter was co-expressed with epidermis marker *Pdpy-7::mCherry*. Their expressions overlapped in epidermis. Bar, 20 μm. (D) tissue-specific expression of *dach-1::SL2::GFP* construct in *dach-1, dpy-7, myo-3*, and *egl-3* promoter all rescued aldicarb-resistance phenotype of *dach-1(ys51)*. **** P<0.0001 (One-way ANOVA, Dunnett’s post-test). (E) *ace-3* fully suppresses the aldicarb resistance of *dach-1*. For (D) and (E), aldicarb sensitivity was measured as the ratio of paralysis at 80 min.

### Pleiotropic effects of *dach-1* mutation on dauer-specfic phenotypic plasticity

We then investigated whether the *dach-1*-dependent altered neurotransmission is important for phenotypic plasticity of dauer in the physiological conditions. Dauer is the hibernation-like stage of *C. elegans* with quiescent movement (Cassada and Russell 1975; Gaglia and Kenyon 2009). Most dauers remain in quiescence, but they can react quickly and move normally after receiving a physical stimulus. Although there was no significant difference in movement and harsh touch sensitivity of *dach-1* dauers compared to wild-type dauers, we found that *dach-1(ys51)* dauers showed relatively decreased quiescence (Figure 4A). When dauers move around 3-dimensional obstacles, they can raise their heads and wave body, which is called nictation (Lee *et al*. 2012; Yang *et al*. 2020). Acetylcholine transmission from IL2 is important for nictation (Lee *et al*. 2012). Also, acetylcholine plays a role in dauer entry (Lee *et al*. 2014). However, neither nictation nor dauer entry was affected by *dach-1* (Figure S5). It shows that *dach-1*-induced synaptic transmission changes do not significantly affect the movement of dauer in laboratory culture conditions.

**Figure 4.**
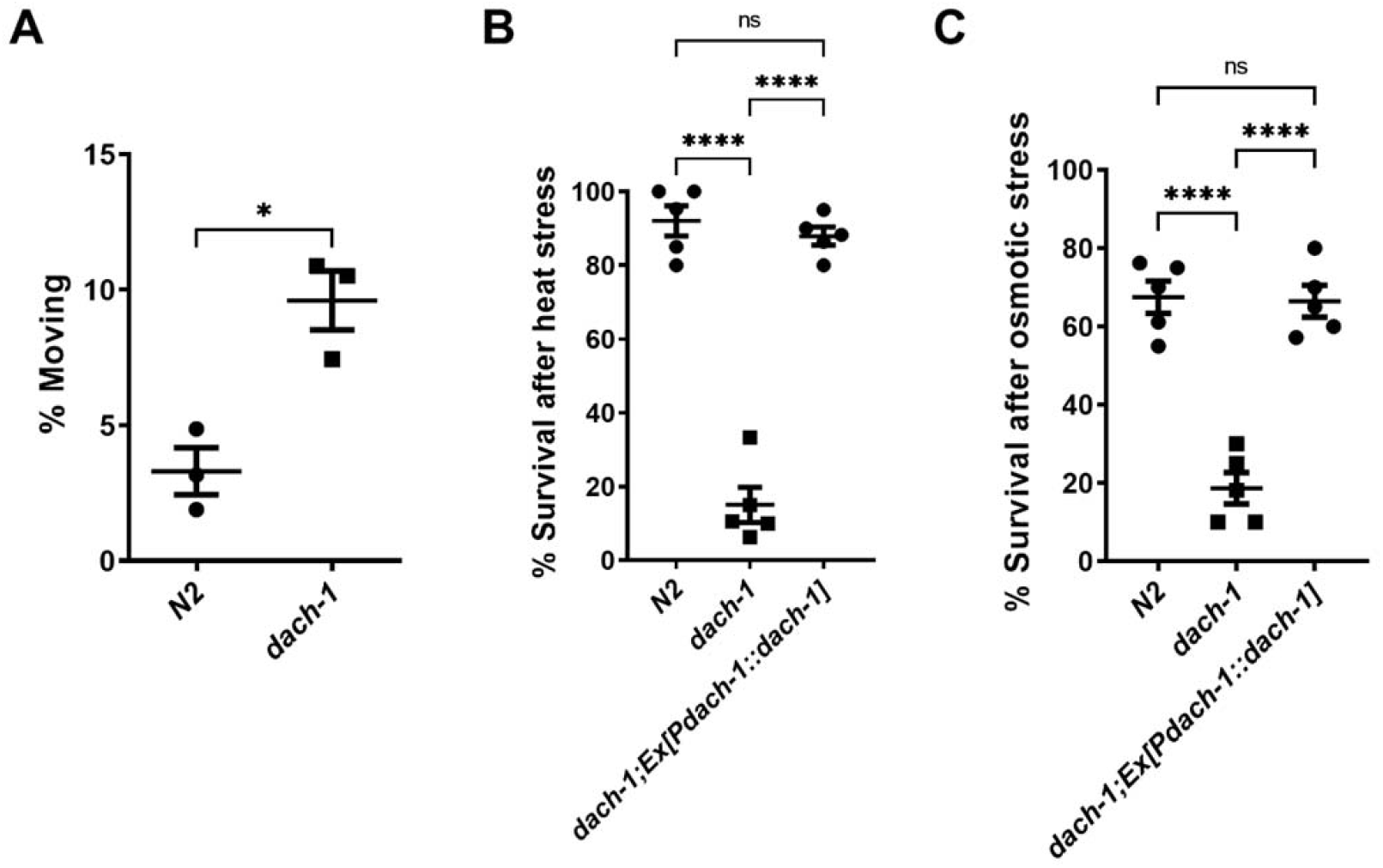
*dach-1* plays a role in quiscence and stress-resistance in dauer. (A) *dach-1* dauer showed increased spontaneous movement. * P<0.05 (Two-tailed t-test). (B and C) *dach-1* dauer is sensitive to heat stress at 37°C for 4 h (B) and high osmotic stress at 1500 mM NaCl for 20 h (C). The sensitivity of *dach-1* to heat stress and osmotic stress was rescued by the expression of *Pdach-1::dach-1::SL2::GFP*. ****P<0.0001 (One-way ANOVA, Dunnett’s post-test)

Another important characteristics of dauer is stress resistance to harsh environments (Golden and Riddle 1984; Narbonne and Roy 2009). Previous studies in *C. elegans* have shown that stress resistance can be regulated by neuronal signaling (Kim and Jin 2015). Interestingly, *dach-1(ys51)* dauers were highly susceptible to heat shock and high osmotic stress (Figures 4B and C). It suggests that the dauer-specific induction of *dach-1* is crucial for the stress response of dauer in harsh conditions. In previous *C. elegans* studies, little was known about the importance of acetylcholine in stress resistance. However, like other aldicarb-resistant mutants, it is possible that *dach-1* may be involved in overall synaptic transmission. Therefore, dauer-specific synaptic plasticity may have modulated stress resistance. However, since acetylcholine neurotransmission and stress resistance are distinct phenotypes, we tend to conclude that *dach-1* has a pleiotropic effect on dauer-specific phenotypes. Overall, we revealed that the function of *dach-1* is important for dauer-specific phenotypic plasticity, especially in stressful environments.

## Discussion

In this study, we reported a previously unreported aldicarb resistance gene specific for dauer. Previously identified aldicarb resistance genes in the adult stage are in most cases genes expressed in neurons, whereas *dach-1* is a gene expressed in the epidermis. It suggests that dauer-specific aldicarb resistance is regulated by a mechanism different from that in adults. Consistently, most aldicarb-resistant mutants at the adult stage had a mild or severe locomotion phenotype, whereas in the case of *dach-1*, no significant difference was found in general locomotion. We discovered that the *dach-1* dauer exhibited less lethargus, a sleep-like behavior, than the wild-type dauer. Since acetylcholine-depleted mutants such as *unc-17* and *cha-1* are more lethargic, the high rate of spontaneous movement with reduced neurotransmission in the *dach-1* dauer is probably not due to cholinergic transmission. Peptidergic signaling in RIS neurons induces dauer sleep-like behavior, similar to sleep-like behavior in other developmental stages (Wu *et al*. 2018). Therefore, the decreased lethargus of *dach-1* is likely due to decreased neurotransmission of RIS or other sleep-inducing neurons.

We found that *dach-1* not only regulates synaptic transmission of dauer, but also plays a pleiotropic effect in stress resistance of dauer. Dauer must survive long without eating. The increased neurotransmission in the dauer seems paradoxical because dauer must use its energy resources efficiently. The stress hypersensitivity of the *dach-1* mutant raises the possibility that the use of energy resources at the synapse was compensated for the survival in stressful environments. In the future, the detailed mechanism underlying the pleiotropic effect may be further explored by examining the direct relationship between the regulation of synaptic activity and stress resistance.

In this study we conducted forward genetics and discovered that *cyp-34A4* mutant dauer was resistant to aldicarb. It is well established that cytochrome P450 is involved in xenobiotic detoxification (Anzenbacher and Anzenbacherova 2001). However, DACH-1 would not have played a role in detoxification of absorbed aldicarb in the body. If that were the case, *dach-1* mutants would be more sensitive to aldicarb instead of resistant compared to wild type. Another role of cytochrome P450 is to participate in the metabolism of cholesterol or polyunsaturated fatty acids to produce intercellular signaling molecules including steroid hormones or epoxyeicosatrienoic acids (Eets) (Gerisch *et al*. 2001; Konkel and Schunck 2011; Spector and Kim 2015). Since cholesterol seems to be not involved in dauer-specific aldicarb sensitivity, *dach-1*-dependent metabolites from polyunsaturated fatty acids may be involved in synaptic transmission. Molecules such as EETs generated through cytochrome P450 metabolism can convey autocrine or paracrine signaling through TRPV channels or ion channels in surrounding cells (Spector and Kim 2015). Hence, the epidermis induction of *dach-1* at the dauer stage identified in our study is likely to play a role in paracrine signaling. Based on the anatomy of *C. elegans*, molecules secreted from the epidermis can be easily transmitted to other neurons and muscles via the pseudocoelom. Also, in *C. elegans* studies, it has been well established that signals from the epidermis regulate glia growth, which in turn regulates the synaptic position of neurons (Shao *et al*. 2013). Therefore, we speculate that the role of the epidermis regulated through *dach-1* may be similar to that of glia or supporting cells in the mammalian brain. Several cytochrome P450s are also expressed in glial cells and are known to be involved in brain inflammation and brain tumors. It would be interesting if the mammalian cytochrome P450 in glia also plays a role in synaptic transmission.

A recent study using *C. elegans* discovered that the transcription of intestinal *dach-1* is regulated by serotonin or dopamine, and the putative signaling molecules generated by *cyp-34A4* regulate the unfolded protein response in the epidermis (Joshi *et al*. 2021). Our results that expression of *dach-1* in the epidermis regulates synaptic transmission suggests that a feedback loop exists in the function of nervous system and the non-neuronal expression of *dach-1*. Another possibility is that *dach-1* may play different roles in distinct tissues at different stages of development. The study of the neuron-to-non-neuronal signaling system involving cytochrome P450 is expected to be of interest in the future. Genes regulating the formation of dauer have been extensively studied through random mutagenesis screening (Fielenbach and Antebi 2008), but studies on genes that change physiology after becoming dauer have been rarely performed. Through random genetic screening, our study found a single gene that regulates developmental plasticity that occurs only at specific stages of development. It suggests a new function of cytochrome P450 in the nervous system during development.

---

**Figure S1.**
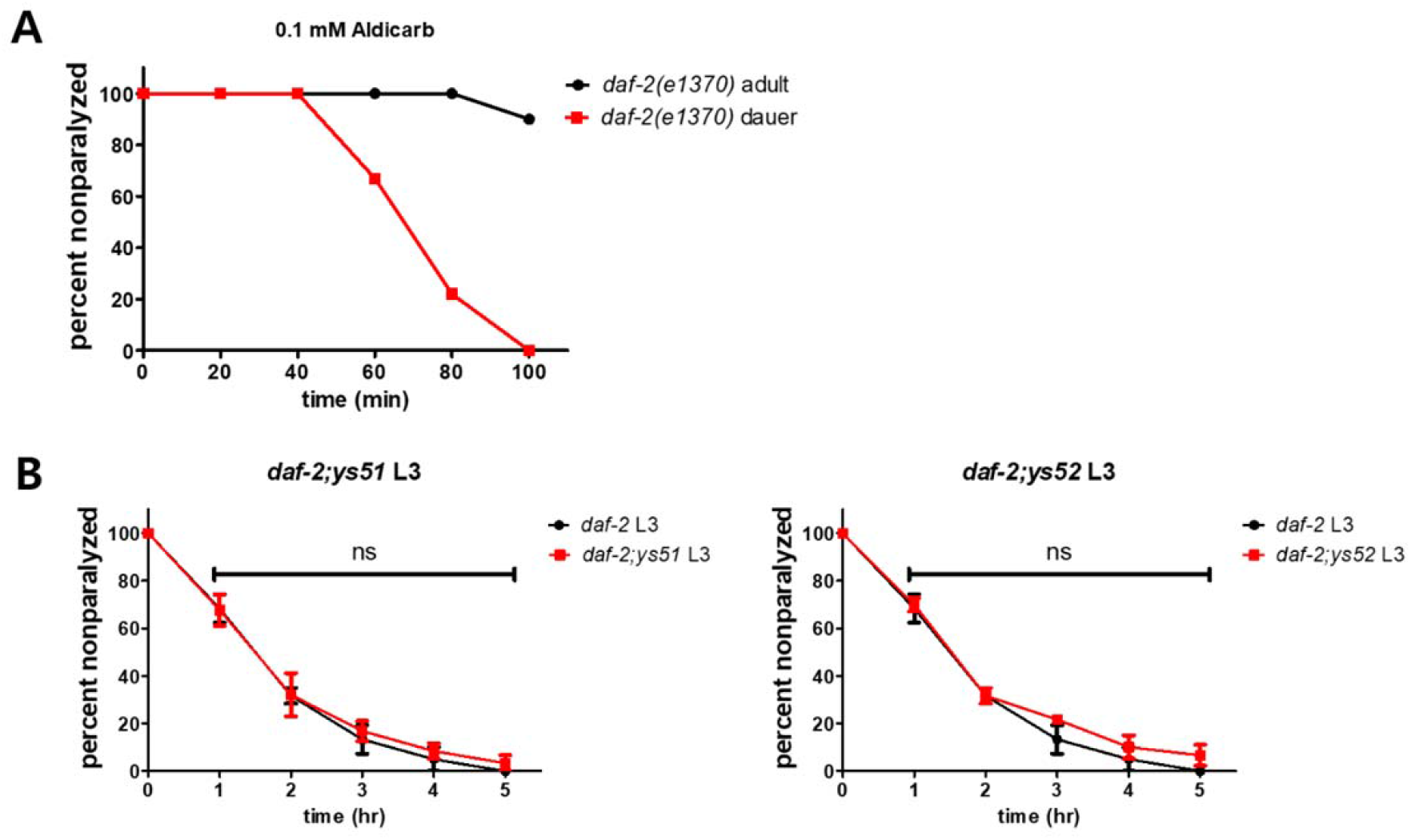
Dauer-specific aldicarb-resistant phenotypes. (A) *daf-2* mutant dauer was sensitive to aldicarb as well. (B) *daf-2;ys51* and *daf-2;ys52* L3 larvae showed similar aldicarb-sensitivity (to 1 mM aldicarb plate) with *daf-2* L3 larvae.

**Figure S2.**
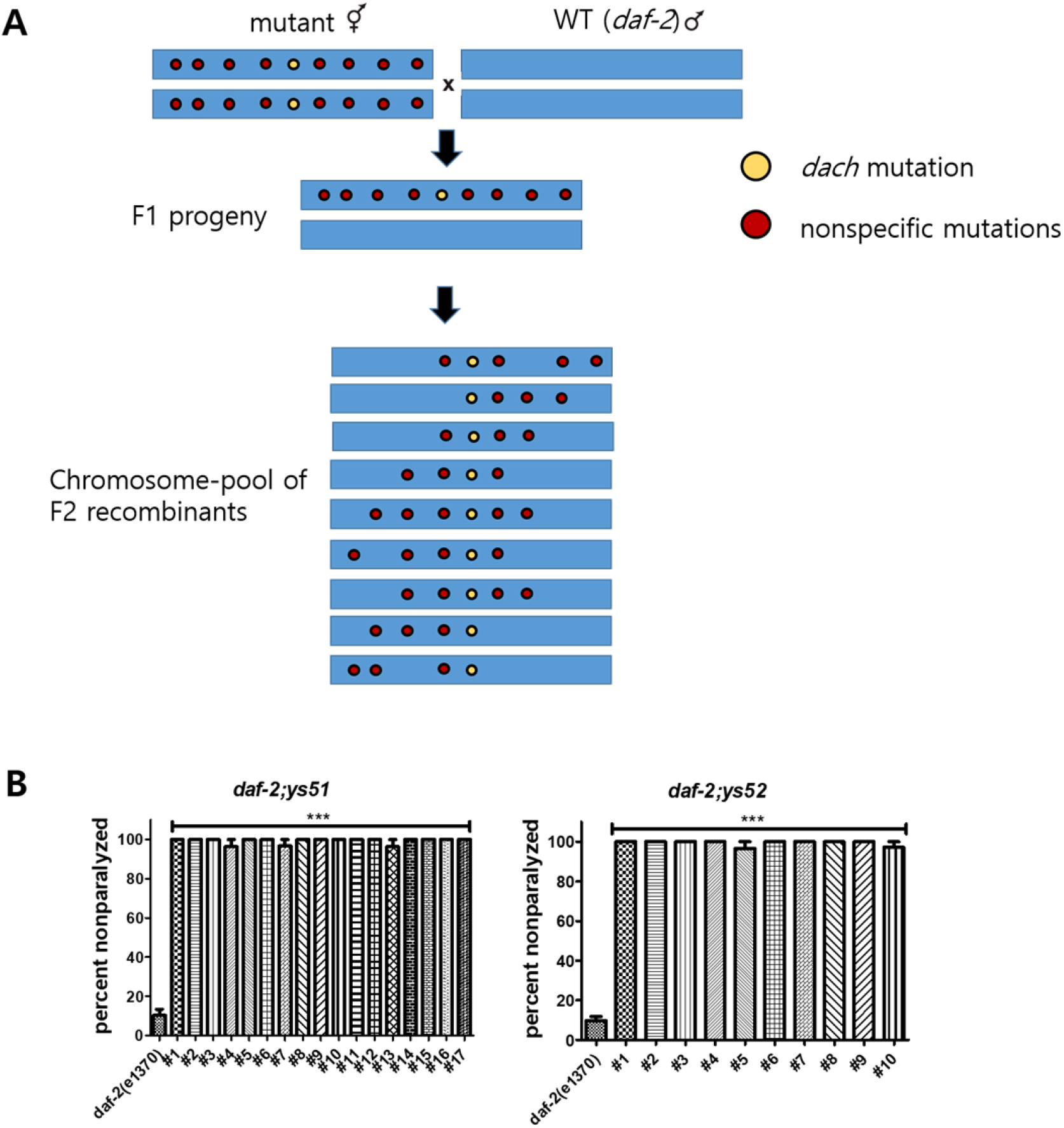
Whole genome sequencing of *dach-1* mutant. (A) Experimental scheme of F2 pooling sequencing strategy. (B) Every independent F2 homozygous was resistant to aldicarb. *** P<0.001 (One-way ANOVA, Dunnett’s post-test).

**Figure S3.**
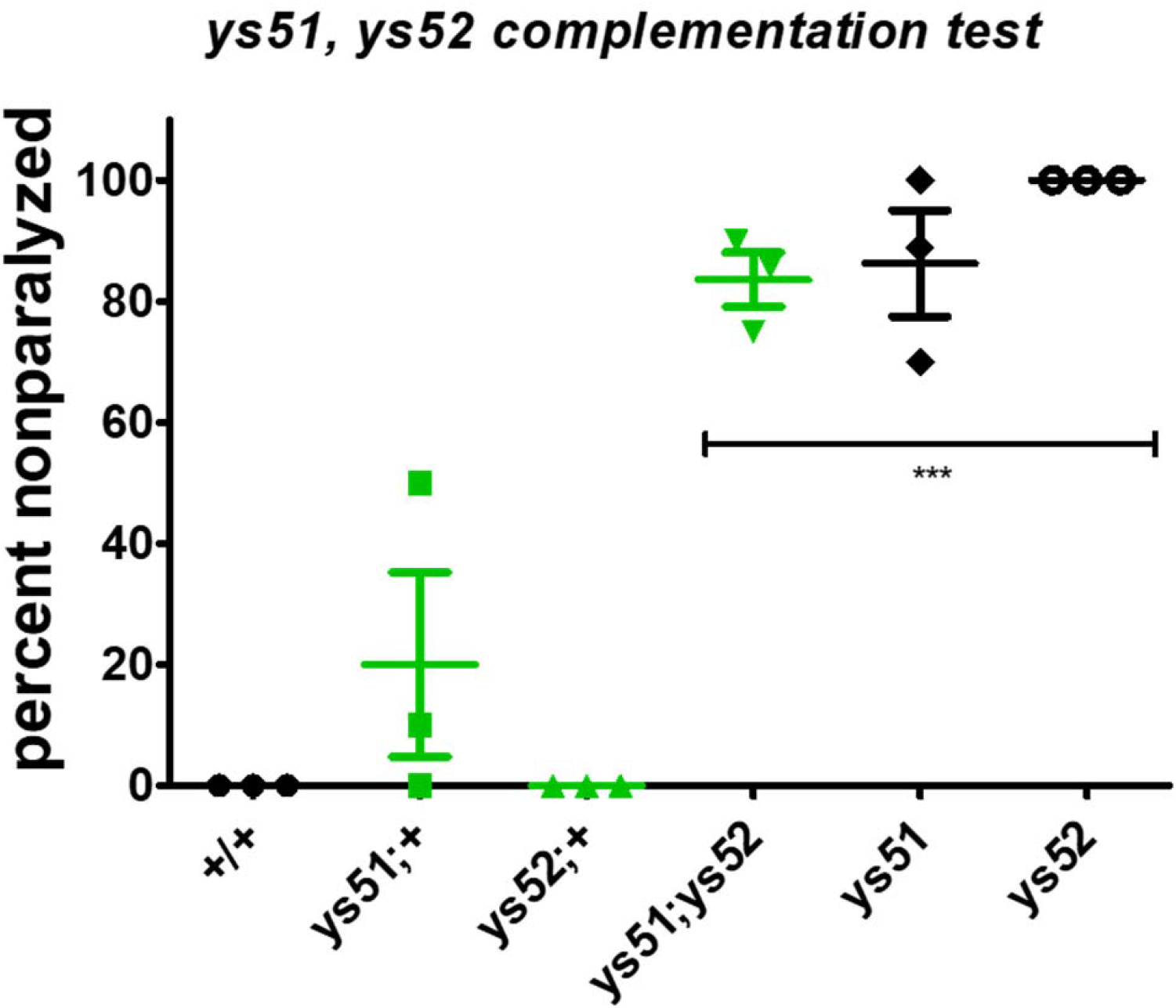
Complementation test of *ys51* and *ys52*. They failed to complement each other, indicating that they share the same mutation. *** P<0.001 (One-way ANOVA, Dunnett’s post-test). Aldicarb sensitivity was measured as the ratio of paralysis at 80 min.

**Figure S4.**
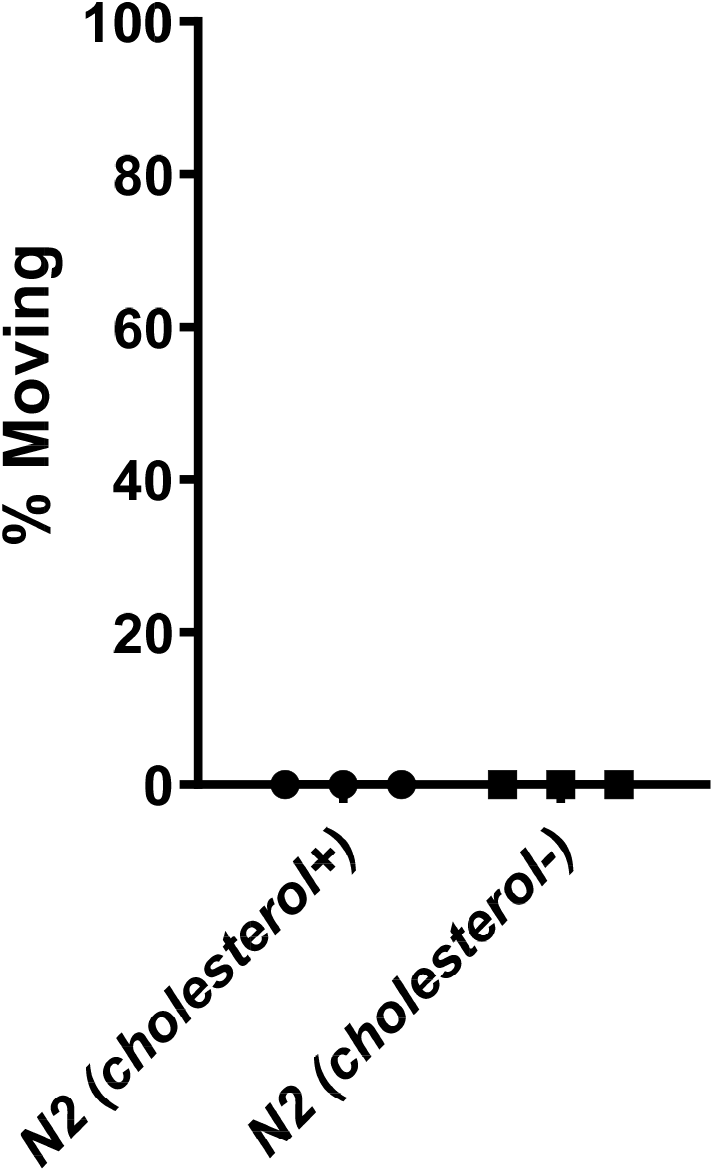
Cholesterol depletion does not influence 0.1 mM aldicarb hypersensitivity at dauer. Aldicarb sensitivity was measured as the ratio of paralysis at 80 min.

**Figure S5.**
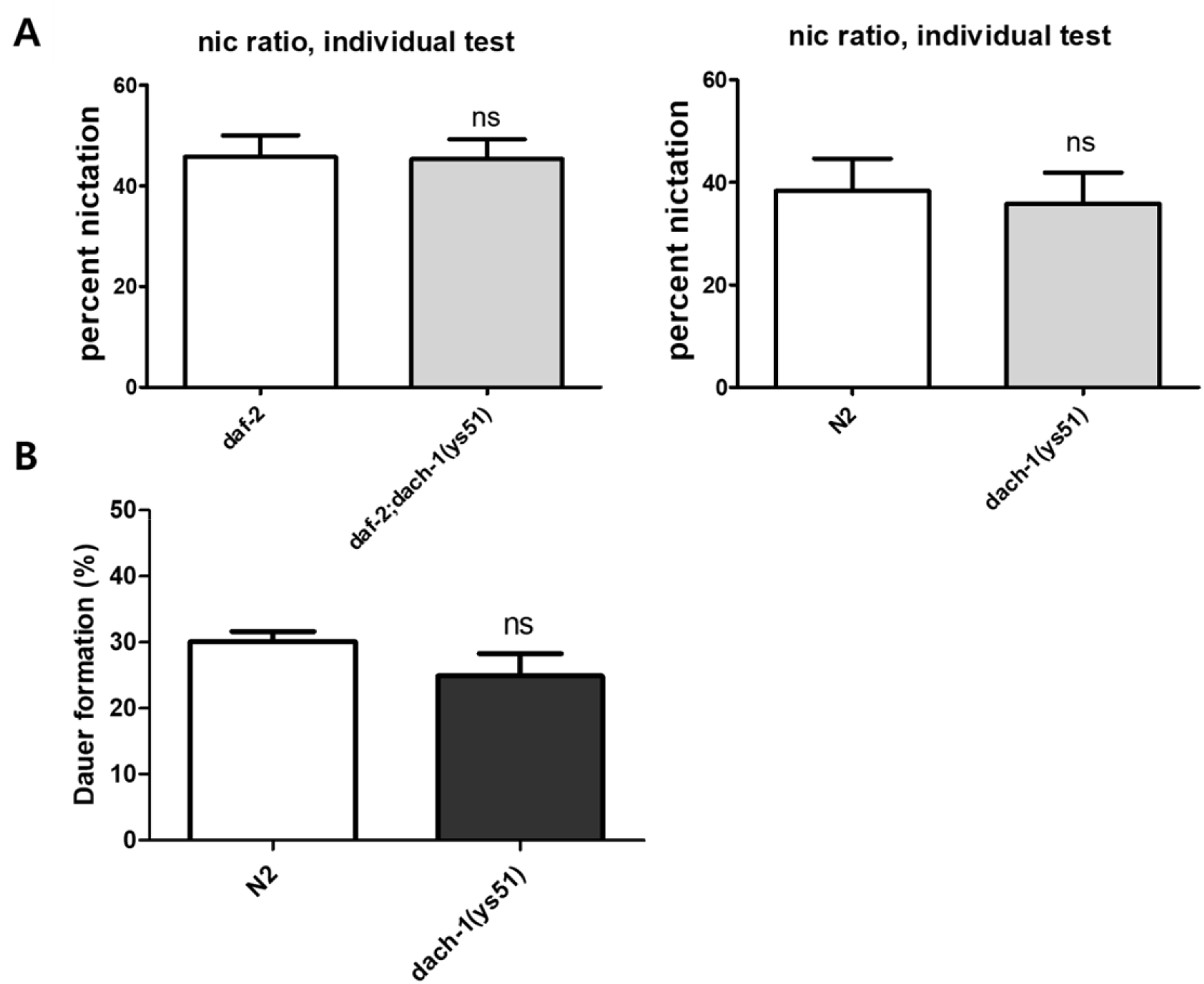
Nictation behavior and dauer formation in *dach-1*. (A) *dach-1* was not defective in nictation, a dauer-specific dispersal behavior. (B) *dach-1* showed normal dauer formation compared to wild type.

## Author contributions

S.S, M.C, and J.L. established the concept of the study. S.S. and M.C. performed the random mutagenesis and genetic analysis. S.S. performed the genetic mapping, the molecular biology experiments, the nictation and lethargus assay. M.C. performed the stress resistance assay. D.S.L. drew the schematic images. J.S. analyzed the whole-genome sequencing data. S.S, M.C, and J.L. analyzed the data and wrote the manuscript. S.S and M.C contributed equally to this work.

## Acknowledgement

*rrf-3(pk1426), daf-2(e1370)*, and *ace-3(dc2)* mutant strains were kindly provided by the *Caenorhabditis* Genetics Center. We thank Dr. Jun Kim, Joost Veroeks, Hyunsoo Yim, Jiwon Do, Chungseok Oh, and Jooyoung Kim for their help with mutant screening and sequencing analysis (Seoul National University). Samsung Basic Science Foundation supported this work.

